# Three-fold Quantum Leap in Butterfly Nectar Knowledge: A Comprehensive Study Revolutionizing Conservation Strategies

**DOI:** 10.1101/2024.09.27.615477

**Authors:** Susan C. Dunlap

## Abstract

This study addresses a critical gap in butterfly conservation: the lack of comprehensive data on nectar plants preferred by US butterflies. The research presented has dramatically improved our understanding, increasing knowledge of nectar plant genera used by butterflies from representing just 25% of all species to an impressive 90%, a more than three-fold or 260% improvement in species representation. This substantial expansion in data has significant implications for conservation strategies and will shift our approach to butterfly habitat restoration. A seven-year longitudinal study of adult butterfly nectaring behavior resulted in over 12,000 detailed records of nectar plant usage by US butterflies. This curated dataset provides a valuable resource with significant implications for all levels of effort directed at butterfly conservation.

Recent recommendations for monarch habitat restoration suggest a balanced mix of host and nectar plants. However, the implementation of this guideline has been hampered by a lack of specific data on monarchs’ preferred nectar plants. New research addresses this gap, providing necessary information to refine plant selection for butterfly habitats. This study also gives cause to modify the current approach of creating generalized pollinator habitats, suggesting that focusing on plants preferred by long-distance migrating butterflies like monarchs may effectively support a broad range of pollinators.

## Introduction

Butterflies play a crucial role in our ecosystems, serving as pollinators and indicators of environmental health. Currently, butterfly populations face numerous threats such as from habitat loss, climate change, and pesticide use, underscoring the urgent need for effective conservation strategies. Critical to butterfly survival is the availability of appropriate nectar plants. Yet, despite their recognized importance, information about these plants has been notably lacking in specificity.

The primary aim of this study is to address gaps in the understanding of butterfly nectar preferences and provide an accessible comprehensive, data-driven resource for land managers, conservationists, and gardeners. To accomplish this, a goal was set to document actual observed nectaring behaviors, providing evidence of butterfly-plant interactions. This goal was met by assembling over 58,000 images and then extracting data from them to compile 12,000+ records of butterfly-plant interactions and document the nectar-feeding behavior of all US butterflies.

Prior to this study, the understanding of nectar plant genera used by butterflies represented only about 25% of all US butterfly species. This significant knowledge gap hampered conservation efforts and limited the effectiveness of habitat restoration projects. New research has dramatically improved this situation, expanding our knowledge to cover 90% of all US butterfly species. This quantum leap in understanding transforms our ability to develop targeted, effective conservation strategies.

A review of existing gardening resources revealed a preponderance of generic lists labeling plants as “butterfly-friendly” without detailing which insect they attract. Numerous taxa on specialized sites listed the adult food as “nectar” or unknown, while other sites listed too few plants to effectively address nation wide conservation needs. The lack of specificity poses significant challenges for those attempting to create targeted butterfly habitats, as gardeners and conservationists often find themselves uncertain about which plants would best support local butterfly populations — an uncertainty that frequently may result in a lack of effective action or expose them to misinformation about the specific nectar plants that appeal to local fauna. Additionally, the habitat development goals of landowners and government agencies may be impeded by the product availability of a limited plant palette.

There has been a growing focus on insect conservation, leading to a surge in efforts to create butterfly habitats in gardens, public lands, and agricultural settings. Despite the considerable resources devoted to preservation, these conservation efforts have been hampered by the lack of specific butterfly-plant relationships. This gap in the conservation literature underscores the need for comprehensive information on the nectar preferences^1^ of US butterflies, particularly those of conservation concern such as monarchs. The lack of comprehensive, genus-specific nectar plant data extends beyond monarchs to encompass all nectar-feeding US butterflies. Thus, allocation of resources to plants that may not effectively support the target insect population is likely quite high.

This study addresses these issues head-on, providing a wealth of new data that dramatically improves our understanding of butterfly-plant interactions. By increasing our knowledge from 25% to 90% of all species, a robust foundation for more effective, targeted conservation efforts has been created. This substantial improvement enables conservationists, land managers, and gardeners to make informed decisions about plant selection, potentially increasing the success rate of butterfly conservation projects and habitat restoration efforts.

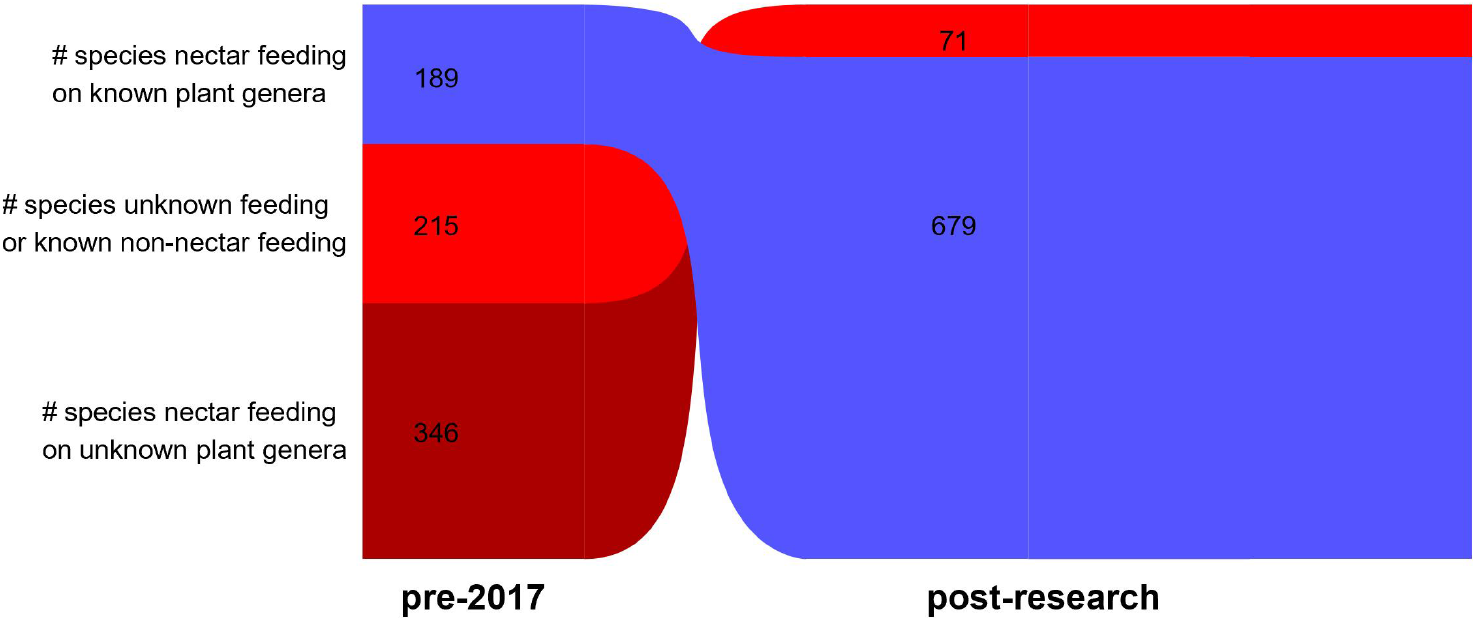
Flower Nectar Data for All 750 US Butterfly Species

The current knowledge gap hampers our capacity to develop tailored conservation strategies for various butterfly species, including those which have poorly documented or unknown nectar preferences. It may be relevant to consider the possibility that butterflies prefer a narrow range of nectar plants, mimicking their frequently tight alignment with a narrow range of host plants.

The absence of a centralized, rigorous database of butterfly-plant interactions represents a significant worldwide obstacle in the field of lepidopteran conservation. Without such a resource, it becomes challenging to make informed decisions about habitat restoration, garden planning, and conservation prioritization. Addressing this problem is crucial for the future success of butterfly conservation.

A recently adopted recommendation in butterfly conservation is the implementation of a balanced mix of host and nectar plants in habitat restoration^2^. While this balanced approach is wise, its effectiveness is compromised by the lack of information about the nectar plants specifically beneficial to each species.

This void in the literature hampers the conservation effectiveness of a guideline that can prove its utility in a variety of conservation efforts when the voids in nectar plant data are overcome. There is a current trend to create habitats beneficial to a broad range of pollinators; this trend is likely to dilute the effectiveness of an effort intended to target a specific species of conservation concern. Parallel to this is a tendency to state a general need for “nectar plants,” without indicating botanical specificity, thus using a term which may expose those tasked to implement a nectar plant mandate to seek suboptimal taxa for a specific conservation project. Current practices often emphasize the selection of plants based on their blooming season; focusing on this attribute can lead to the inclusion of plants that may not be preferred or even utilized by the target butterfly species.

These extant approaches highlight the need for insect-specific information on nectar plant preferences. By addressing the limitations in our current knowledge, and remedying them, we can enhance the effectiveness of conservation efforts and provide more urgently needed support for butterfly populations.

## Methodology

A library of over 58,000 images from numerous publicly available online resources and vetted aggregators^3^ was assembled and then a standardized observation protocol was employed to document butterfly nectaring events. The images recorded detailed information of butterfly-plant interactions, including a visual record of a butterfly species, its nectaring behavior, floral details, and the environmental conditions. The image database included a complete set of images for US butterflies, as needed to discern male and female species, and auxiliary images of native wild flora so-as to develop familiarity with each taxon’s distinguishing characteristics. Numerous online resources were used to compile and cross-check the US-inclusive butterfly taxa^4^. The auxiliary images supported a working library of over 32,000 images of novel butterfly-plant interactions, each one explicitly recording a butterfly’s active feeding behavior. Central to the approach was to apply knowledge acquired during a ten-year study to develop a patented plant identification method^5^, thus eliminating much of the ambiguity that has plagued previous studies; the method combined novel and traditional taxonomic approaches with a careful review of distinctive floral traits. Images of unknown butterflies and plants were identified using online image recognition software^6^ and benefited from the accumulated knowledge acquired as the project advanced. The process was constantly refined to become more discriminating in selecting references and sources and learning from identification errors — sorting through thistles, yellow Aster blossoms, and the Eupatorium group was particularly challenging. The image collection process improved over time, primarily by pre-screening images with insufficient identification data. Ambiguities were resolved or abandoned by referencing vetted Lepidoptera and botanical resources^7^.

The butterfly images were collected in distinct folders to affirm continuity and accuracy. When distinct floral characteristics were noted for an individual butterfly-plant image, a specific moniker was assigned to the image. When only partial details were recorded, an effort to identify the plant species was made by diving deep into botanical and horticultural literature and pushing image identification resources to their current limits. An image was abandoned if a genus-level identification was not achieved; thus, images identified above the genus level were not added to the master database. Plants identified to genus or below were placed in the appropriate plant family and genus folders. Assembling the images in this manner improved the accuracy of the data, as the commonality between images in each folder served to affirm the validity of genera assignments that had been made.

Custom information that applied to each image was then added to each record, including butterfly species and common name, plant family, genus, and, where possible, plant species. Location data was not typically recorded; further research is needed to add this information to the resource. Initially, when the vetted aggregator community was in its infancy, images were gathered from a broad range of online resources; these images infrequently included specific data. This approach was later abandoned in favor of collecting images from vetted aggregators. The image software’s export features^8^ allowed a direct export of the image data to be managed in a spreadsheet. Excel analytic tools were used to process, examine, and interpret the vast amount of information and to identify patterns in butterfly-plant interactions. Additional data columns were added to the master database including plant type, USDA plant native and naturalized status^9^, host plant data^10^, USDA hardiness planting zones^11^, butterfly formal name^12^, dominant genera species counts, and butterfly range in the US^13^. Numerous online resources were used to cross-check the butterfly taxa^14^. More recently, AI^15^ has been used to acquire plant details, such as naturalizing and self-sow behaviors, information that was cross-checked for accuracy. Numerous subdivisions of the master database were created to facilitate a deeper examination of an insect or plant of interest, such as monarchs, dominant plants, plant families, and Asclepias. The data was sorted, reviewed, edited, and studied for multiple attributes, commonalities, and distinctions. Each action revealed compelling and consequential data. Such analysis is ongoing.

This comprehensive longitudinal data collection covered a wide geographic range, included a review of images collected between 1959 and 2024, and required seven years to create. This extended timeframe was crucial for capturing the full spectrum of butterfly behavior. All US butterfly species were considered. The breadth of the study ensured that data representative of diverse ecosystems and climate zones was captured. The inclusive approach may set the research apart from previous studies. By including all species, a goal was met to provide a complete picture of butterfly nectar preferences across all US butterfly fauna. This approach allowed for consistency in data gathering. Throughout the study, rigorous data validation and quality control measures were employed, and the process to secure consistency in the data gathering and achieve productive results was continuously refined.

The comprehensive nature of this study allowed us to dramatically expand our understanding of butterfly nectar preferences. Prior to this research, only about 25% of US butterfly species had documented nectar plant genera associations. Our goal was to significantly increase this coverage, aiming for a more complete picture of butterfly-plant interactions across the United States.

## Findings

The research as of 24Sept24 yielded a wealth of new information about the nectar preferences of US butterflies, dramatically improving our understanding from 25% to 90% coverage of all species. In total, 12,334 detailed records of butterfly nectaring behavior were compiled. This dataset represents the most comprehensive collection of US butterfly-plant interactions to date and provided a robust foundation for analyzing patterns in each butterfly’s feeding preferences. Key findings include:

1. Of the extant 750 US butterfly species, 679 (90.53%) are nectar-feeding.
2. Undocumented feeding behaviors in several butterfly species were discovered:
  - Novel data covered 100 of 142 butterflies whose adult feeding behavior was previously unknown, a 71% decrease in unknown behavior.
  - 400 of 453 butterflies previously designated as “nectar feeding” without specific plant species assignments now have detailed nectar plant preferences, an 88% decrease in “nectar feeding” designation.
3. 664 genera in 124 plant families servicing the needs of all nectar-feeding US butterflies were identified.
  - 446 of these genera contain species that can be grown in nearly every state in the continental US.
  - 86 Dominant Genera, each with a minimum of 30 records, appeal to 640 (94.25%) US nectar-feeding butterfly species.
  - 92% of these genera contain plant forms that will either self-sow or naturalize in a variety of settings.
4. “Keystone” Genera:
  - 10 “Keystone” Genera, each with a minimum of 200 records, appeal to 503 (74%) US nectar-feeding butterfly species.
  - 27 “Keystone+” Genera, each with a minimum of 100 records, appeal to 527 (77.6%) US nectar-feeding butterfly species.
5. Monarch butterfly findings:
  - 1,672 records (13.14% of total) are of monarchs.
  - Monarchs were observed nectaring on 232 genera.
  - 21 plant genera were identified as dominant to monarchs, with 95% of these containing taxa that either self-sow or naturalize in various US settings.
6. *Asclepias* (milkweed) findings:
  - 689 records cover *Asclepias*.
  - 188 US butterflies (25% of all US butterflies) have been observed nectaring on *Asclepias*.

These observations highlight the importance of comprehensive, sustained studies in revealing the full spectrum of butterfly behavior. A review of the Dominant Genera shows that *Asteraceae* is the commanding plant family with 40.52% of the records, followed by *Verbenaceae* at 10.94%, and *Lamiaceae* at 6.81%. The “Keystone” Genera embody low plant diversity numbers with an extremely high yield of US butterflies attracted to feed. The “Keystone” Genera (in order) are: *Asclepias, Cirsium, Lantana, Verbena, Salvia, Symphyotrichum, Bidens, Zinnia, Trifolium*, and *Eriogonum*. The “Keystone+” group adds (in order): *Buddleja, Conoclinium, Verbesina, Taraxacum, Echinacea, Centaurea, Solidago, Liatris, Chromolaena, Monarda, Rudbeckia, Phyla, Gaillardia, Heliotropium, Pentas, Vernonia*, and *Helianthus*. These genera contain 1082 species that grow in the United States. The diverse growth forms of these taxa include: 496 annuals, 114 biennials, and 2585 perennials^16^.

It is noteworthy that *Asclepias*, the monarch’s host plant, dominates the favorite plant list for both the general butterfly population and monarchs themselves. Very few individual taxa have such broad appeal and consequential conservation effectiveness. It is reasonable to consider that the sustained conservation effort mounted on behalf of monarchs has contributed to the general appeal of *Asclepias* to a great many US butterflies. As such, it serves as an affirmation of what dedicated and sustained conservation efforts can achieve. Applying the same effort to other plants which have conservation appeal may yield similar benefits for other beneficial insects.

In all, monarchs were observed nectaring on 232 genera. The Dominant Genera of particular importance to monarchs are (in order): *Cirsium, Lantana, Verbena, Bidens, Asclepias, Salvia, Zinnia, Conoclinium, Solidago, Buddleja, Helianthus, Eupatorium, Symphyotrichum, Liatris, Echinacea, Trifolium, Taraxacum, Pentas, Verbesina, Tithonia*, and *Coreopsis*. Overall, these taxa are comprised of 75 perennials, 4 biennials, 11 annuals, and 11 shrubs. Such information has immediate practical applications for the creation and management of monarch conservation. It is worth noting that perennials and annuals top the list of each Dominant group examined — monarch favorites and favorites of all US nectar-feeding butterflies. The data shows that adding *Eupatorium, Tithonia*, and *Coreopsis* to a “Keystone and Keystone Genera+” conservation plan will insert all the monarch preferred genera into that plan. The diverse plant form totals would then include: 518 annuals, 117 biennials, 2728 perennials^17^.

Cultivating the monarchs’ favored nectar plants shows great benefit to them and appeals to 574 (84.6%) nectar-feeding US butterflies. If the monarch’s preferred nectar plants become more widely available, by self-sowing and naturalizing in gardens and fields, they may appeal to a broader subset of US butterflies than the current data suggests. Thus, the successful appeal of *Asclepias* can be replicated to expand the broad appeal of these monarch-friendly nectar plants to then feed other US butterflies. Planting the monarch’s statistically desirable perennials, biennials, and annuals that naturalize or self-sow may provide great benefit to the whole butterfly population. Further examination will show which taxa to add to the monarch-favored plant list that will then benefit the remaining 12 (15.3%) US nectar-feeding butterflies without having to compromise monarch restoration strategies. Further study will likely demonstrate that the monarch’s favorite nectar plants appeal to a considerable number of pollinating insects, including bees, moths, and birds. It is reasonable to consider that a plant’s naturalizing behavior has drawn the favorable attention of many pollinating fauna; attention that may have been noted by the plant itself.

Additionally, tracking the feeding behavior of specific butterflies may enable researchers to deduce an image’s location. Further study may reveal significant regional variations in butterfly nectar preferences. The current data contains all available data for some insects, particularly those with a limited range and those whose nectar-feeding behavior had been unknown. For other insects, the counts may increase but not impact the current number of genera. Given the continent-wide range of monarchs, they may have evolved a preference for naturalizing taxa, as they gravitated to a frequently encountered floral form with recognized olfactory appeal — attracted by a familiar field-grown meal.

These findings represent a quantum leap in our understanding of butterfly nectar preferences. By increasing our knowledge of all species, we have created a robust foundation for more effective, targeted conservation efforts. This dramatic improvement enables us to make more informed decisions about habitat restoration and management, potentially increasing the success rate of butterfly conservation projects.

## Discussion

The dramatic improvement in our understanding of butterfly nectar preferences, from 25% to 90% coverage of all species, has far-reaching implications for butterfly conservation efforts. This quantum leap in knowledge provides a scientific foundation for managing habitats for a wide range of US butterfly species.

Foremost, the research has significant implications for monarch butterfly conservation efforts. The detailed data on monarch nectar preferences, now part of a much more comprehensive dataset, provide a scientific foundation for more effective habitat creation and management. Conservation organizations and land managers can now make more informed decisions about plant selection for monarch way stations, potentially increasing their survival rates during migration. Furthermore, by selecting locally naturalizing taxa from the list of preferred genera for insects of conservation concern, they can disperse seeds to initiate the development of a self-sustaining habitat. This targeted approach to habitat restoration could play a crucial role in reversing the alarming decline in butterfly populations, including monarchs^18^.

Numerous butterflies are highly specialized in their preference of host plants, typified by the fact that *Asclepias* is the Monarch’s solitary host plant. This study suggests that they are also highly specialized nectar feeders that we have been treating as nectar generalists. We can now become as specialized as the butterflies by planting their preferred nectar plants. Monarchs are also known to prefer dense plantings of their host plant^19^, especially when planted in relatively small clusters at optimal distances apart, thus responsive to their propensity and instinct to migrate. It may also be found that monarchs favor optimally spaced, dense plantings of their preferred nectar plants. Monarch’s olfactory skills serve them well during migration, as it is known they can detect blooming host plants for “several hundred meters,^20^” but what is their detection range when a preferred nectar plant is in bloom (and *Asclepias* is not)? It may be possible to determine the detectable range of an olfactory plume relative to a narrow set of nectar plants. Future plantings guided by the wisdom of such findings can align with the monarch’s migratory strengths.

Beyond monarchs, these findings may have broad implications for the conservation of all US butterfly species, and perhaps other insects around the world. The comprehensive nature of the study, encompassing 90% of US nectar-feeding species (up from the previous 25%), allows for the development of more nuanced and species-specific conservation strategies. This is particularly important for rare or endangered butterfly species, whose specific nectar requirements may have been previously unknown or poorly understood. A list of locally adapted host plant options may prove to be just as useful for conservation planning as what has been found to be true of nectar plants.

In the realm of habitat restoration and management, the research challenges some existing practices while reinforcing others. The identification of “Keystone” Genera that support multiple butterfly species provides a powerful tool for maximizing the impact of conservation efforts, especially those with limited resources, with the caveat that future studies may indicate that narrow nectar plant preferences apply to some butterflies. The current findings underscore the need for locally adapted approaches rather than one-size-fits-all solutions. Such approaches can help landowners create habitat that serves them well in the long term, as they are tutored to avoid invasive species within an otherwise desirable genus. Additionally, locally adapted approaches can explore the narrow nectar-plant hypotheses with ease.

The significant role played by perennials, biennials, and annuals cannot be overemphasized, as the majority of plants attractive to butterflies either naturalize or self-sow: an impressive 92% of the favored taxa for all nectar-feeding US butterflies and 95% of the favored taxa for monarchs. These percentages underscore the advantage of disseminating nectar-plant information to all who are concerned about butterfly conservation. The findings indicate that use of a plant palette comprised of naturalizing species will prove to be a highly effective approach to long-term conservation of key butterfly species. With care, conservationists can broadcast these favored seeds all over the land.

From a scientific perspective, this study significantly advances the field of pollinator ecology. The discovery of previously undocumented feeding behaviors and the detailed mapping of nectar preferences across 90% of species, opens new avenues for research into butterfly evolution, ecology, and behavior.

This research also has implications for climate change studies and conservation planning. By providing a comprehensive baseline of current butterfly-plant interactions, this data can be used to track and predict changes in these relationships as climate patterns shift. This information could be crucial for developing adaptive conservation strategies worldwide in the face of environmental change.

For gardeners and urban planners, this research offers a scientifically grounded guide for creating more effective butterfly gardens and pollinator-friendly urban spaces. By providing specific information on self- sowing and self-sustaining nectar plants for different butterfly species, the findings can help increase the success rate of butterfly gardening efforts, potentially leading to more robust urban and suburban butterfly populations.

The implications of this research extend beyond the immediate field of butterfly conservation. By providing a deeper understanding of the intricate relationships between butterflies and their nectar sources for 90% of species, this study contributes to broader efforts in biodiversity conservation, sustainable land management, and ecological research. The methodology employed and the resulting database may set a new standard for comprehensive ecological studies.

While the potential for future study is broad, a few particular areas of conservation may improve by attending to issues revealed by these findings. Primary conservation literature promoted the theme “all hands on deck^21^” which was then adopted by numerous agencies^22^ to drive home the need to attend to the diminishing and threatened Western and Eastern migrating monarch populations. Those agencies and nonprofits will benefit from the information gathered in this database, as they continue to serve the large land managers that form the basis of their outreach. Gardeners, among others, constitute an audience that would reply to the conservation appeal proffered by a wise use of this data. Offering them an extensive list of available garden plants known to be favored by 94% of US nectar-feeding butterflies including monarchs would potentially put thousands of acres of land into play, as 60% of US land mass is under the care of private landowners^23^. For gardeners and urban planners, this research offers a scientifically grounded guide for creating more effective butterfly gardens and pollinator-friendly urban spaces.

Focused use of the self-sowing and self-sustaining nectar plant findings can help increase the success rate of butterfly gardening efforts, potentially leading to more robust urban and rural butterfly populations.

In addition to providing vital information for the existing nationally focused conservation community, there are structures in place that could expand exponentially. One such infrastructure — comprised of the thousands of growers, garden centers, and seed purveyors that serve the gardening community — can become co-collaborators to help restore the US monarch populations and other endangered insects, especially if they maintain a pledge to void the use of harmful chemicals and provide naturalizing forms of nectar plant taxa. This nectar plant data makes it possible to appeal to this broad and diverse audience of known butterfly enthusiasts.

The implications of this research may extend beyond the immediate field of butterfly conservation. By providing a deeper understanding of the intricate relationships between butterflies and their nectar sources, this study contributes to broader efforts in biodiversity conservation, sustainable land management, and ecological research. The methodology employed and the resulting database may set a new standard for comprehensive ecological studies. This approach could serve as a model for similar large-scale investigations of other pollinator groups or plant-animal interactions.

Of special note is the need for further study to vindicate choosing plants that are known to conserve one species, that may indeed support the conservation of many other pollinators as well. We can knowingly support monarchs, an endangered, beloved, and extraordinary world-class migrating species, while gathering more information about the conservation needs of other pollinators and endangered insects.

Future studies may underscore the importance of tailoring conservation efforts to local ecological conditions and butterfly populations, rather than relying on one-size-fits-all approaches. The data can be explored for these variations by isolating records for specific regional butterfly species. For example, taxa that may naturalize too aggressively in one region of the country may be perfect nectar additions to another. Noting the specific behavior of each species within a favored genus could prove to be invaluably useful information for gardeners and other conservationists. The study identified several plant species that appear to be disproportionately important as nectar sources for multiple butterfly species. These “Keystone” Genera could be prioritized in conservation efforts to maximize benefits for butterfly populations.

Given that human recording of these nectaring events was essential to the creation of this body of work, it must be noted with caution that the images gathered may only partially reflect or embody an individual butterfly’s feeding preferences. Further study is required to 1) identify more plants to the species level, 2) discern if a butterfly has specific, rather than broad, nectar plant preferences, 3) prepare a list of Dominant Genera for other insects of conservation concern, 4) reveal whether plants that appeal to some US butterflies contain fewer self-sowing taxa so are less inclined to naturalize, and 4) illuminate regional variations in butterfly nectar preferences. This database may prove to be an essential asset to address these studies. The representativeness of this data collection necessitates consideration of the gap between reality and what can be collected. It is important to acknowledge that some of the image data may reflect nectar choices made under duress. However, despite these potential limitations, this database and its curation are currently the most comprehensive available. As such, it serves as an essential asset for addressing these concerns and advancing the understanding of the subject.

The findings reveal intriguing patterns in nectar plant preferences among different butterflies; whether these choices are by familiarity or convenience is not revealed in the data. However, there are certain patterns revealed by studying the preferred plants and their propagation behavior. These patterns may prove useful in both predicting nectaring behavior and planning conservation efforts. Much can be gained by further study of the data, providing a glimpse into butterflies’ preferred dining flora. While some butterflies showed a high degree of specialization in their nectar sources, others demonstrated remarkable flexibility; the data provides insights on how to distinguish between these feeding preferences.

## Conclusion

This comprehensive study of nectar preferences among US butterflies addresses a critical gap in butterfly conservation knowledge, dramatically improving our understanding from 25% to 90% coverage of all species. By identifying key “Keystone” Genera and documenting previously unknown feeding behaviors, this research provides a solid foundation for more effective and targeted conservation efforts.

The findings emphasize the importance of naturalizing and self-sowing plants in butterfly habitats, particularly for species of conservation concern like the monarch butterfly. This work not only contributes significantly to the field of pollinator ecology but also offers practical guidance for conservationists, gardeners, and land managers. As we face ongoing challenges in biodiversity conservation, this quantum leap in understanding provides valuable insights that can inform adaptive strategies in the face of environmental change.

The correlation between humans and insects highlights a co-dependency that can be used to guide the conservation actions we take on behalf of insects. Following their lead to cultivate naturalizing taxa on their behalf can be done with much greater precision now that we understand the nectar preferences of 90% of US butterfly species, and regardless of our capacity to record each feeding event. With care, conservationists can broadcast these favored seeds all over the land, supporting not just a few, but the vast majority of US butterfly species. The data shows a strong overlap between most US butterflies and a relatively compact list of nectar plants. This overlap can make future conservation efforts more expansive, inclusive, efficient, and simpler.

I am grateful to Steve Port for his sustained support and guidance in the development of this paper and extend special thanks to Jennifer Thieme of Monarch Joint Venture and Changzhi Ai PhD.

## Footnotes

1 I explicitly acknowledge the difficulty of defining a butterfly’s nectar preference, and am using the term to reflect my examination of over 58,000 publicaly available images that span over 25 years of US butterfly nectaring events.

2 The Midwest Association of Fish and Wildlife Agencies. 2023 Update to the Mid-America Monarch Conservation Strategy 2018-2038.

3 https://www.inaturalist.org, https://www.gbif.org, https://images.google.com

4 https://www.butterfliesofamerica.com, https://www.butterfliesandmoths.org, https://www.gbif.org, https://bugguide.net

5 https://ppubs.uspto.gov/dirsearch-public/patents/html/8301389?source=USPAT&requestToken=eyJzdWIiOiJhZjJhZDM4MC1jMDc0LTQzZjktOWZhYi0xZTczMTYzZTYzYjYiLCJ2ZXIiOiI5YmFhNmU3Ny04YzlhLTQ1YmUtYjJhOC0zNmZhZTdlMjM5ZmIiLCJleHAiOjB9 https://ppubs.uspto.gov/dirsearch-public/patents/html/8577616?source=USPAT&requestToken=eyJzdWIiOiJhZjJhZDM4MC1jMDc0LTQzZjktOWZhYi0xZTczMTYzZTYzYjYiLCJ2ZXIiOiI5YmFhNmU3Ny04YzlhLTQ1YmUtYjJhOC0zNmZhZTdlMjM5ZmIiLCJleHAiOjB9

6 https://images.google.com

7 https://www.butterfliesofamerica.com, https://www.butterfliesandmoths.org, https://www.gbif.org, https://bugguide.net, https://www.worldplants.de/world-plants-complete-list/complete-plant-list

8 Adobe Lightroom Classic

9 https://plants.usda.gov/home

10 https://data.nhm.ac.uk/dataset/hosts, https://openai.com/chatgpt, https://claude.ai

11 https://planthardiness.ars.usda.gov/, https://www.missouribotanicalgarden.org, https://www.gardenia.net/, https://en.wikipedia.org/wiki, https://pfaf.org

12 https://www.gbif.org

13 https://www.gbif.org, https://www.butterfliesandmoths.org

14 https://www.butterfliesofamerica.com, https://www.butterfliesandmoths.org, https://www.gbif.org

15 https://claude.ai

16 https://plants.usda.gov/home; these numbers are fungible depending on the plant zone where a taxon is grown. The plant form total also includes 51 shrubs, 2 vines, and 3 trees.

17 The plant form total includes 59 shrubs.

18 Brice X. Semmens, et al., 2016, Quasi-extinction risk and population targets for the Eastern, migratory population of monarch butterflies (Danaus plexippus)

19 Anna Syke Bruce, et al., 2021. Landscape- and local-level variables affect monarchs in Midwest grasslands

20 Blackiston et al. 2011; Grant et al. 2018

21 W.E. Thogmartin et al., 2017, Restoring monarch butterfly habitat in the Midwestern U.S.: “All hands on deck.”

22 2023 Update to the Mid-America Monarch Conservation Strategy 2018-2038, The Midwest Association of Fish and Wildlife Agencies

23 www.summitpost.org/public-and-private-land-percentages-by-us-states/186111

## Notes

### Competing Interest Statement

The authors have declared no competing interest.

